# Evolution of the sex-determination gene *Doublesex* within the termite lineage

**DOI:** 10.1101/2024.05.07.592865

**Authors:** Kokuto Fujiwara, Satoshi Miyazaki, Kiyoto Maekawa

## Abstract

The molecular mechanism of sex determination has long been considered conserved in insects. However, recent studies of hemimetabolous insects have challenged this notion. One notable example is termites. In *Reticulitermes speratus*, a homolog of sex determination gene, *Doublesex* (*RsDsx*), exhibits characteristics that are distinct from those of other insects, including sister-group cockroaches. It comprises a single exon, contains only doublesex/mab-3 DNA-binding domain (DM) but lacks a conserved oligomerization domain (OD), and exhibits transcriptional activity only in males. To investigate whether these characteristics are widespread within the termite lineage, we identified *Dsx* homologs in three different families. The loss of the conserved OD sequences was observed in all termite species examined, whereas the number of exons and expression patterns between sexes varied among families. Particularly, distinctive differences in *Dsx* were found in species from the Archotermopsidae and Kalotermitidae, both of which have a linear caste developmental pathway. Our findings indicate that diversification of *Dsx* structure and expression patterns may have contributed to ecological diversification, such as caste developmental pathways, within the termite lineage.

## 1. Introduction

Clarifying the molecular mechanisms underlying sex determination and differentiation is a crucial challenge in biology. The sex of insects is determined by diverse genetic factors that are unique to each lineage. Examples, include X chromosome dosage in the fruit fly *Drosophila melanogaster* (Erickson and Quintero, 2007), a sex determination factor on the sex chromosome in the Mediterranean fruit fly *Ceratitis capitata* (Meccariello et al., 2019), heterozygosity of the *complementary sex determiner* in the honeybee *Apis mellifera* (Beye et al., 2003), and the maternal effect genomic imprinting sex determination system in the parasitoid wasp *Nasonia vitripennis* (Van De Zande and Verhulst, 2014). Sex information is transmitted through multiple genes that constitute the sex determination pathway and ultimately leading to sexual differentiation.

Advances in sequencing technology helped identify gene homologs that constitute the sex determination pathway in many insect species (Geuverink and Beukeboom, 2014). Specifically, a sex determination gene, *Doublesex* (*Dsx*), encoding the transcription factor has been analyzed for its structure, expression patterns, and functions in many species. *Dsx* is a member of the doublesex- and mab-3 related transcription factors (DMRTs) family and possesses two conserved domains: doublesex/mab-3 DNA-binding domain (DM) and oligomerization domain (OD) (An et al., 1996; Matson and Zarkower, 2012; Raymond et al., 1998). Sex-specific isoforms of *Dsx* are produced through regulation by the upstream splicing factor (e.g. *transformer* and *P-element somatic inhibitor*) (Hoshijima et al., 1991; Suzuki et al., 2008). Formation of sex-specific traits is regulated by sex-specific isoforms of *Dsx*. These characteristics were conserved in holometabolous insects (Price et al., 2015; Shukla and Nagaraju, 2010; Verhulst and Van de zande, 2015). However, distinct characteristics have been reported in species other than holometabolous insects. For example, splicing variants of *Dsx* exist, but all isoforms are expressed regardless of sexes in the louse *Pediculus humanus* and the silverleaf whitefly *Bemisia tabaci* (Guo et al., 2018; Wexler et al., 2019). Furthermore, *Dsx* regulates differentiation only in males but not in females in the German cockroach *Blattella germanica* and the firebrat *Thermobia domestica* (Chikami et al., 2022; Wexler et al., 2019). Notably, *Dsx* of the termite *Reticulitermes speratus* (*RsDsx*) consists of a single exon with a DM domain (loss of the conserved OD sequences) and is transcribed in males but not in females (Miyazaki et al., 2021). *Cryptocercus* cockroaches are a sister group of termites, and sex-specific *Dsx* isoforms of *C. punctulatus* (containing both DM and OD domains) have been identified (Miyazaki et al., 2021). Consequently, termites are one of the most important groups to understand the evolution of *Dsx*-dependent sex-determination pathway in insects. In termites, information on *Dsx* (gene structure, number of exons, expression pattern, and predicted target genes) has been clarified in *R. speratus* (Fujiwara et al., 2024; Miyazaki et al., 2021), whereas fragmentary information exists for other species.

In this study, we searched for *Dsx* homologs in three termites belonging to three families: *Zootermopsis nevadensis* (Archotermopsidae), *Gryptotermes satsumensis* (Kalotermitidae), and *Odontotermes formosanus* (Termiridae) (Figure 1). Additionally, we attempted to verify whether *Dsx* homologs in *Hodotermopsis sjostedti* (Archotermopsidae) consist of a single exon with no available genomic information. Finally, the expression patterns of *Dsx* were compared between males and females. Based on these results, we discuss the commonalities and diversity of *Dsx*, and the evolutionary process of *Dsx*-related ecological traits within the termite lineage.

**Figure 1.**
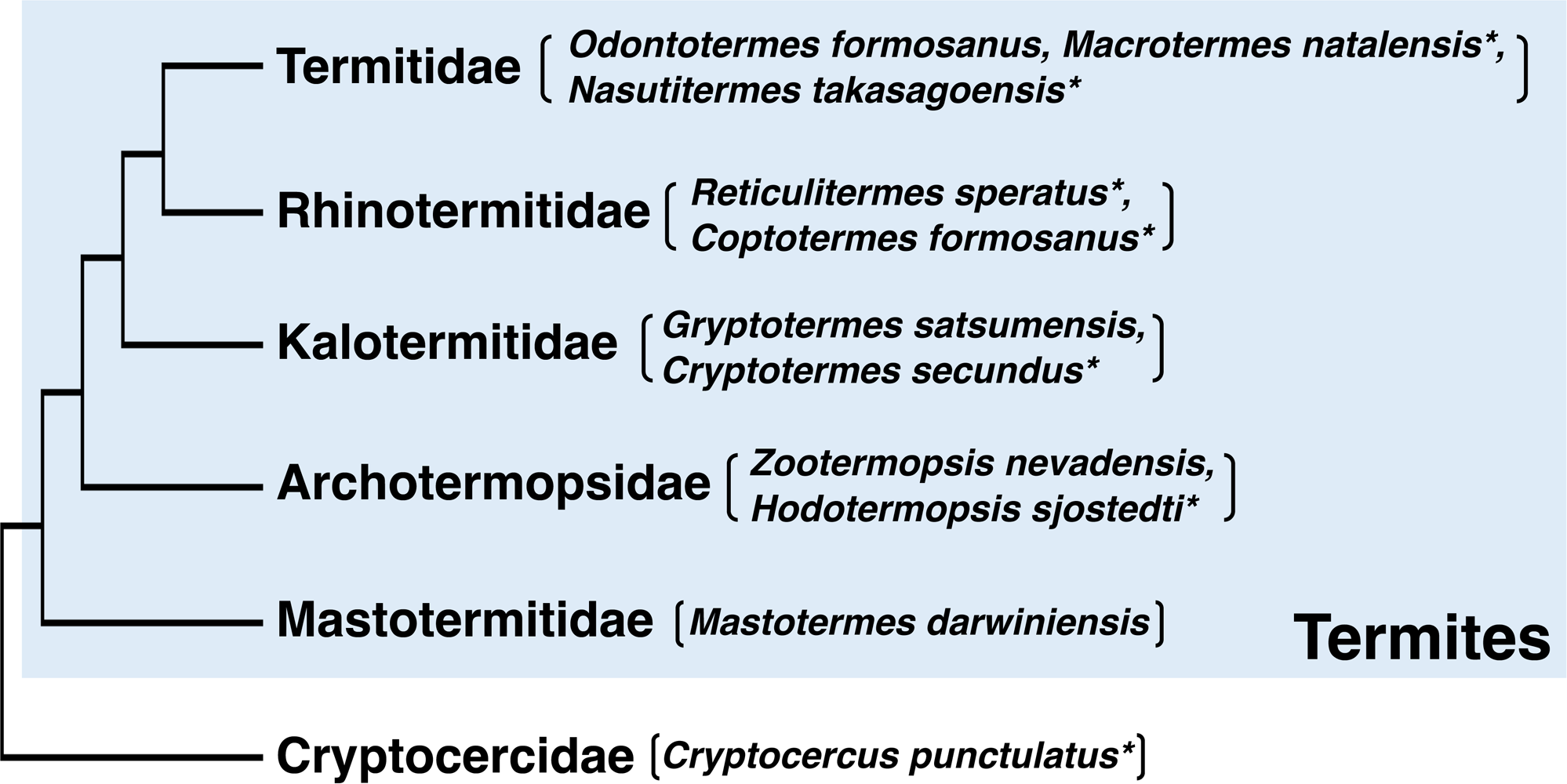
Phylogenetic relationship between termites and sister group *Cryptocerucus* cockroaches. A simplified phylogenetic tree shows the relationships among termite families and sister group cockroach family Cryptocercidae (including only *Cryptocercus* spp.). The species used in this study are indicated in parentheses. Asterisks denote species for which the *Dsx* homologs were reported in the previous study (Miyazaki et al. 2021).

## 2. Material and methods

### 2.1. Termites

Seven colonies of *Z. nevadensis* were collected from Kawanishi, Hyogo Prefecture, Japan, in April 2023. Three colonies of *H. sjostedti* and one colony of *G. satsumensis* were collected from Yakushima, Kagoshima Prefecture, Japan, in May 2023. Three colonies of *O. formosanus* were collected from Naha, Okinawa Prefecture, Japan, in April 2022. The colonies were maintained in plastic cases at room temperature. Note that in species belonging to the families Archotermopsidae and Kalotermitidae, individuals designated as working immatures within the colony are typically referred to as pseudoergates (false-workers); however, in this study, they are simply referred to as workers.

Three colonies of *Z. nevadensis* were used to collect of workers and soldiers, and the remaining four colonies were used to collect of winged imagoes (alates). Newly molted alates from different colonies were used to establish incipient colonies according to previous study (Maekawa et al., 2012). Female and male individuals were paired in a 60 mm plastic petri dish containing nest wood chips. All dishes were kept in an incubator at 25 degrees in constant darkness. Approximately 2.5 months after colony establishment, five incipient colonies were used for the collection of primary reproductives. Three colonies of *H. sjostedti* were used to collect workers and soldiers, and one colony was used to collect alates. Three colonies of *O. formosanus* were used to collect workers and alates. A colony of *G. satsumensis* was used to collect last-instar nymphs.

### 2.2. RNA-seq analysis of *G. satsumensis*

Total RNA was extracted from the whole body of each individual (male and female nymphs) using ReliaPrep RNA Tissue Miniprep Systems (Promega, Madison, WI, USA). The quality and quantity of the extracted RNA were evaluated using a NanoVue spectrophotometer (GE Healthcare Bio-Sciences, Uppsala, Sweden) and a Qubit 2 fluorometer (Thermo Fisher Scientific, Waltham, MA, USA). The total RNA extracted from male and female nymphs was mixed in equal amounts. Directional mRNA library preparation and sequencing (total 82,108,272 reads) were performed by Novogene Co., Ltd. (Tianjin, China) using NovaSeq at 150 bp paired-end (PE) reads. The adapter sequences, low quality (< q30), and short sequences (< 50 bp) were removed using fastp v0.20.1 (Chen et al., 2018). After quality control, *de novo* assembly sequences were constructed using Trinityrnaseq v2.15.0 (Grabherr et al., 2011). All reads will be deposited in the DDBJ Sequence Read Archive (DRA) database.

### 2.3. *De novo* assembly analysis of *Z. nevadensis* RNA-seq data

*De novo* assembly analysis was performed using previous RNA-seq data (accession: PRJNA203244; Terrapon et al., 2014). Data obtained from male reproductive castes (both the primary and secondary male reproductives) were used for this analysis. Removal of adapters, low quality, and short sequences, and *de novo* assembly analysis were performed using the methods described above.

### 2.4. Searching for Dsx in G. satsumensis and Z. nevadensis

To identify candidate *Dsx* homologs, blastn was performed against *de novo* assembly sequences obtained in each species using BLAST+ 2.12.0 (Camacho et al., 2009). DM domain sequences of other termites (*Cryptotermes secundus,* and *H. sjostedti*; Miyazaki et al., 2021) were used as queries.

### 2.5. Searching for *Dsx* in *O. formosanus*

Genomic DNA (gDNA) was extracted from the whole bodies of workers (25 individuals) using ISOSPIN Tissue DNA (Nippon Gene, Tokyo, Japan). Primer sequences were designed for the highly conserved regions of *Dsx* sequences in *Macrotermes natalensis* and *Nastitermes takasagoensis* (Miyazaki et al., 2021), both of which belong to the same family as *O. formosanus* (Termitidae) (Table S1). PCR was performed using KOD FX Neo (Toyobo, Osaka, Japan). The obtained product was purified using the QIAquick Gel Extraction Kit (QIAGEN, Tokyo, Japan). After adapting of the overhanging dA, the purified product was subcloned into a pTA2 Vector (Toyobo), and transfected into *Escherichia coli* XL1-blue. The inserted DNA sequences were determined using the BigDye Terminator v3.1 Cycle Sequencing Kit (Thermo Fisher Scientific) with an Applied Biosystems 3500 Series Genetic Analyzer (Thermo Fisher Scientific).

### 2.6. 5□ /3□ Rapid amplification of cDNA ends (RACE)

Total RNA was extracted from the male reproductives of *Z. nevadensis* (whole body of one individual) and *O. formosanus* (head and thorax of one individual) using the ISOGEN □ (Nippon Gene). The extracted RNA was purified using DNase □ (TaKaRa Bio, Shiga, Japan). cDNA was synthesized using a SMARTer RACE cDNA Amplification Kit (Clontech Laboratories, Mountain View, CA, USA). Gene-specific primers were designed using Primer3 Plus (Untergasser et al., 2007) (Table S2). RACE was performed using Advantage 2 Polymerase Mix (Clontech Laboratories). The obtained 5□- and 3□-RACE products were purified using a QIAquick Gel Extraction Kit (QIAGEN), subcloned into a pTAC-2 Vector (Bio Dynamics Laboratory lnc., Tokyo, Japan), and transfected into *E. coli* XL1-blue. The inserted DNA sequences were determined using the QuantumDye Terminator Cycle Sequencing Kit v3.1 (Tomy Digital Biology Co., Ltd., Tokyo, Japan) with an Applied Biosystems 3500 Series Genetic Analyzer (Thermo Fisher Scientific). The obtained cDNA sequences were confirmed to exhibit high similarity to the known *Dsx* homologs using a BLAST search. All sequences will be deposited in the DDBJ/EMBL/GenBank databases.

### 2.7. Molecular phylogenetic analysis of DM domain sequences

A phylogenetic tree of DM domain sequences was constructed according to a previous study (Miyazaki et al., 2021). We used the newly obtained sequences, as well as those previously used for the phylogenetic analysis (nine termites, *C. punctulatus*, *B. germanica*, and *D. melanogaster*) (Table S1). Phylogenetic relationships were inferred using the maximum likelihood (ML) method (bootstrap value of 1000) in MEGA X (Kumar et al., 2018). The most appropriate model was determined using the model selection option implemented in MEGA X, and the K2 + G model was selected.

### 2.8. Analysis of exon structure

The total gDNA of four species (*Z. nevadensis*, *H. sjostedti*, *G. satsumensis*, and *R. speratus*) was extracted from male and female workers using QIAGEN Genomic-tips 20/G (QIAGEN). One individual was used for DNA extraction in the first three species, whereas five individuals were used in *R. speratus* because of their small body size. The amplification of *beta-actin* was performed to confirm the success of gDNA extraction using the primers designed for conserved regions in all examined termites (Table S2). The total RNA of each species was extracted from male and female nymphs (each one individual, whole body) using ISOGEN □ (Nippon Gene) and ReliaPrep RNA Tissue Miniprep Systems (Promega). Total RNA extracted from a male and female nymph was mixed in equal amounts. cDNA was synthesized from the extracted RNA using a High-Capacity cDNA Reverse Transcription Kit (Thermo Fisher Scientific). *Dsx, pros*, and *UTX*-specific primers for each species were designed from the obtained sequences using Primer3 Plus (Untergasser et al., 2007) (Table S2). PCR amplification of the sequences was performed using Ex Premier DNA Polymerase (TaKaRa Bio).

### 2.9. RNA extraction for gene expression analysis

The total RNA of *Z. nevadensis* and *H. sjostedti* was extracted from each caste (reproductives, soldiers, and workers) using ISOGEN □ (Nippon Gene). The total RNA of *Z. nevadensis* was extracted from one individual (head or remaining body part) per sample from three different colonies, and quintuplicate biological samples were prepared for each category. In *H. sjostedti*, the total RNA of the reproductives, soldiers, and workers was extracted from one individual (head or remaining body part) per sample from three different colonies (soldiers and workers) or a single colony (reproductives), and triplicate biological samples were prepared. Extracted total RNAs of both species were purified using DNase I (TaKaRa Bio). Total RNA of *G. satsumensis* was extracted from male and female last instar nymphs using the ReliaPrep RNA Tissue Miniprep Systems (Promega). Total RNA was extracted from one individual (head and thorax or abdomen) per sample from a single colony, and triplicate - quadruplicate biological samples were prepared for each category. Total RNA of *O. formosanus* was extracted from male and female reproductives using the ReliaPrep RNA Tissue Miniprep Systems (Promega). Total RNA was extracted from one individual (head and thorax) per sample from a single colony, and triplicate biological samples were prepared for each category. Total RNA from the abdomen of *O. formosanus* was excluded in analysis because of its low purity. The quality and quantity of extracted RNA were measured using a NanoVue spectrophotometer (GE Healthcare Bio-Sciences) and a Qubit 2 fluorometer (Thermo Fisher Scientific). cDNA was synthesized from the purified RNA using a High-Capacity cDNA Reverse Transcription Kit (Thermo Fisher Scientific).

### 2.10. Real-time quantitative PCR analysis

Gene expression levels of *Dsx* homologs were measured using the Thunderbird SYBR qPCR Mix (Toyobo) and the MiniOpticon Real-time PCR system (Bio-Rad, Hercules, CA, USA) (*G. satsumensis* and *O. formosanus*) or the QuantStudio 3 Real-Time PCR System (Thermo Fisher Scientific) (*Z. nevadensis* and *H. sjostedti*). To determine an internal control gene, according to previous studies (Gao et al., 2020; Masuoka et al., 2015; Oguchi et al., 2022; Terrapon et al., 2014; Xu et al., 2021), the suitability of reference genes was evaluated using GeNorm (Vandesompele et al., 2002) and NormFinder (Andersen et al., 2004) software (Table S3). Gene-specific primers were designed using Primer3Plus (Untergasser et al., 2007) (Table S2). The dot plot graphs of expression levels were drawn using the “ggplot2” R package (Ginestet, 2011). These studies were conducted using R ver. 4.1.2 (available at https://cran.r-project.org/).

### 2.11. Gene expression analysis in *C. secundus*

Gene expression levels of *Dsx* homologs were measured using previous RNA-seq data (accession: PRJNA382129; Harrison et al., 2018). The data of reproductives (queens and kings) were used in this analysis. Adapter, low quality, and short sequences were removed using the methods described above. Transcript expression was quantified using salmon v1.4.0 (Patro et al., 2017). Transcript data from *C. secundus* (GCF_002891405.2: PRJNA432597; Harrison et al., 2018) were used as a reference data set. Transcript expression levels were quantified as transcripts per million (TPM). The dot plot graphs of expression levels were drawn using the methods described above.

### 2.12. Statistical analysis

*Dsx* expression levels were compared among sexes (males and females) and castes (reproductive, workers, and soldiers) using a two-way ANOVA followed by Tukey’s test (p < 0.05) in *Z. nevadensis* and *H. sjostedti*. *Dsx* expression levels were compared between sexes (male and female) using Student’s t-test for *O. formosanus*, *G. satsumensis*, and *C. secundus*. All statistical analyses were performed using Mac Statistical Analysis ver. 2.0 (Esumi, Tokyo, Japan).

## 3. Results

### 3.1. Searching for *Dsx* in termites

In a previous study (Miyazaki et al., 2021), *Dsx* homologs were identified in six termites. Here, we searched for *Dsx* homologs in three termite species belonging to three families (*Z. nevadensis*, *G. satsumensis*, and *O. formosanus*) (Figure 1). We obtained a single sequence in *Z. nevadensis* (*ZnDsx*: 648 bp, 215 aa) (Figure S1) and *O. formosanus* (*OfDsx*: 1,107 bp, 368aa) (Figure S2), and two sequences in *G. satsumensis* (*GsDsx1*: 2379 bp, 792 aa; *GsDsx2*: 1026 bp, 341 aa) (Figure S3-4). All obtained sequences possessed DM domains but lacked OD domains (Figure S1-4). To confirm the orthology of *Dsx* in termites, we performed phylogenetic analysis using the DM domain nucleotide sequences of termites, cockroaches (*C. punctulatus* and *B. germanica*), and *D. melanogaster*. Reportedly, each family of DMRTs (*Dmrt11*, *Dmrt93*, *Dmrt99*, and *Dsx*) forms a distinct clade (Chikami et al., 2022; Miyazaki et al., 2021). The resultant tree showed that all DM domain sequences obtained were clustered within the *Dsx* clade (Figure 2, Table S1). In *Cryptotermes secundus* belonging to the Kalotermitidae, which is the same family as *G. satsumensis*, two *Dsx* homologs have been identified (*CsDsx1* and *CsDsx2*; Miyazaki et al., 2021). Each pair of *Dsx* identified in *G. satsumensis* and *C. secundus* (i.e., *GsDsx1* and *CsDsx1*, *GsDsx2* and *CsDsx2*) were closely related phylogenetically (Figure 2).

**Figure 2.**
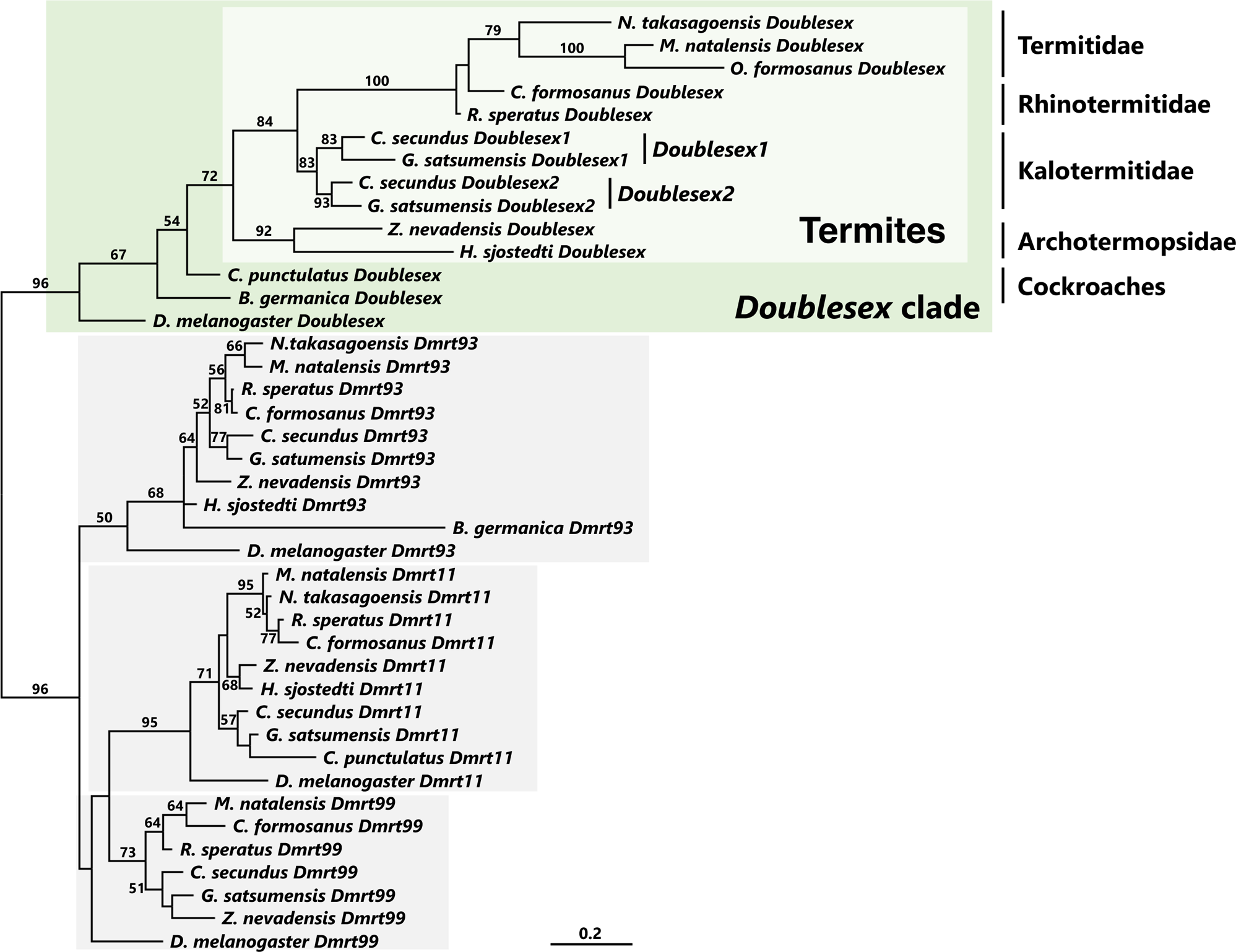
Phylogenetic tree of DM domain sequences in termites and cockroaches. Maximum likelihood tree was constructed based on the DM domain sequences of *Dsx* and *Dmrt* homologs. The numbers above or below branches indicate the level of support in bootstrap analysis (1000 replicates, only above 50% was shown). *Dsx* clade is indicated by green box. Each *Dmrt* clade is indicated by gray box.

### 3.2. PCR amplification using genomic DNA (gDNA) and complementary DNA (cDNA)

*Dsx* homologs consisted of a single exon structure in all termites examined in a previous study (Miyazaki et al., 2021). To clarify the exon structure of the newly identified *Dsx*, we focused on *Dsx* homologs in *Z. nevadensis* and *G. satsumensis*, both of which could be analyzed using a large number of samples. First, using the *Z. nevadensis* genome sequence (Terrapon et al., 2014), we attempted to determine the genomic position of *ZnDsx*. However, no homologous sequences were identified. Next, we investigated the amplification product size of *ZnDsx* using gDNA and cDNA (Figure 3). *ZnDsx* was amplified from male gDNA but not from female gDNA (Figure 4a) using gene-specific primers (Table S2). Product sizes were much larger than those obtained using a mixture of cDNA derived from female and male nymphs (Figure 4a). Using the same method and gene-specific primers (Table S2), we investigated the PCR product sizes for *G. satsumensis*. The same sizes of PCR products were obtained using gDNA and cDNA for both paralogs (Figure 4b-c). The *Dsx* of *R. speratus* (*RsDsx*) consists of a single exon (Miyazaki et al., 2021), and we confirmed that the PCR product sizes using gDNA and cDNA were the same (Figure 4d). To verify whether the difference of product size derived from gDNA and cDNA is commonly observed in the family Archotermopsidae, we focused on the *Dsx* of *H. sjostedti* (*HsDsx*: Miyazaki et al., 2021). Amplification using gene-specific primers (Table S2) was confirmed only in the male gDNAs, and the PCR product sizes using gDNA and cDNA were completely different (Figure 4e).

**Figure 3.**
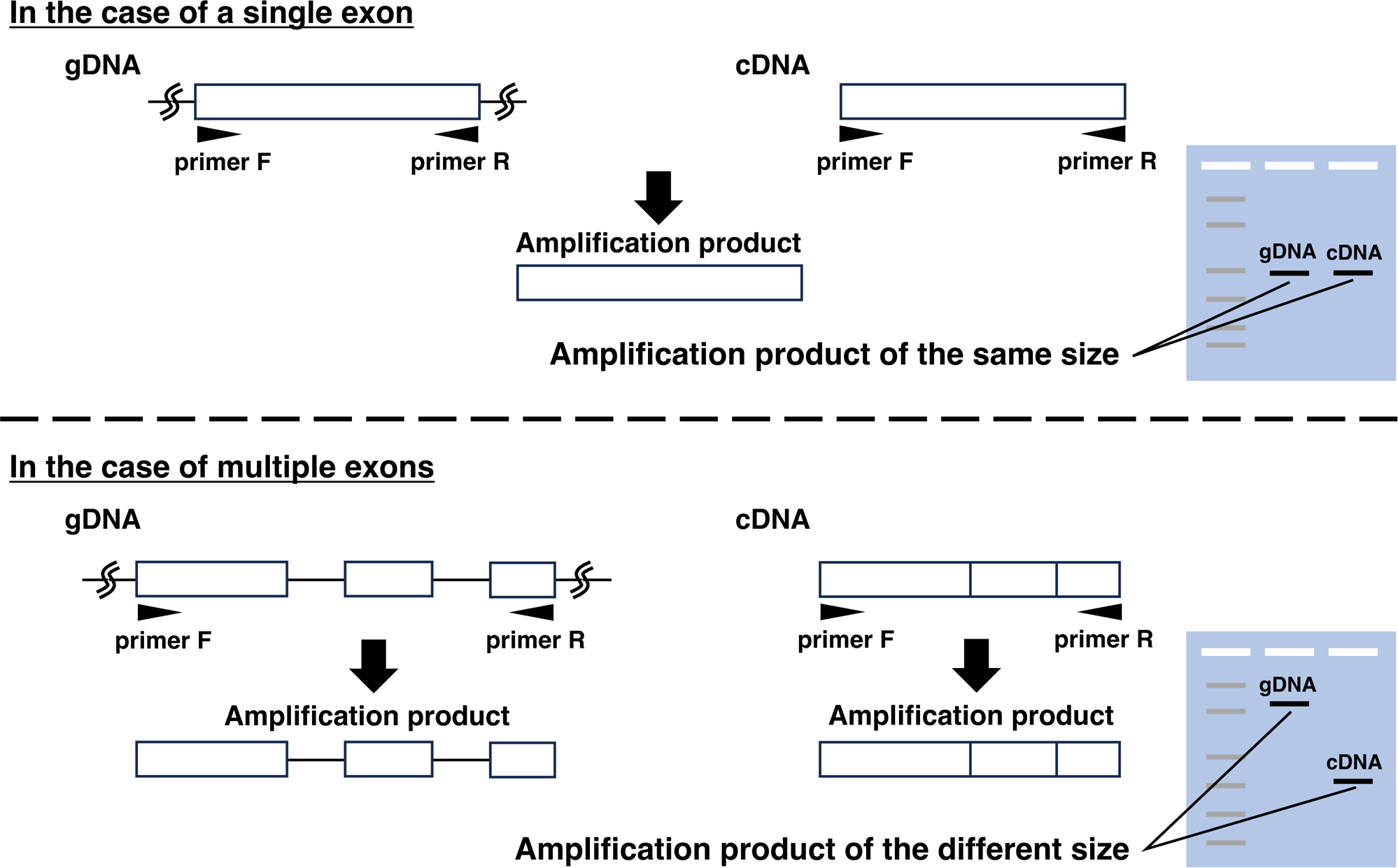
Experimental design for exon structure analysis. In the case of a single exon structure, the amplification product sizes using gDNA and cDNA are the same. In the case of multiple exon structures, the amplification product size using gDNA is larger than that using cDNA due to the presence of intron sequences.

**Figure 4.**
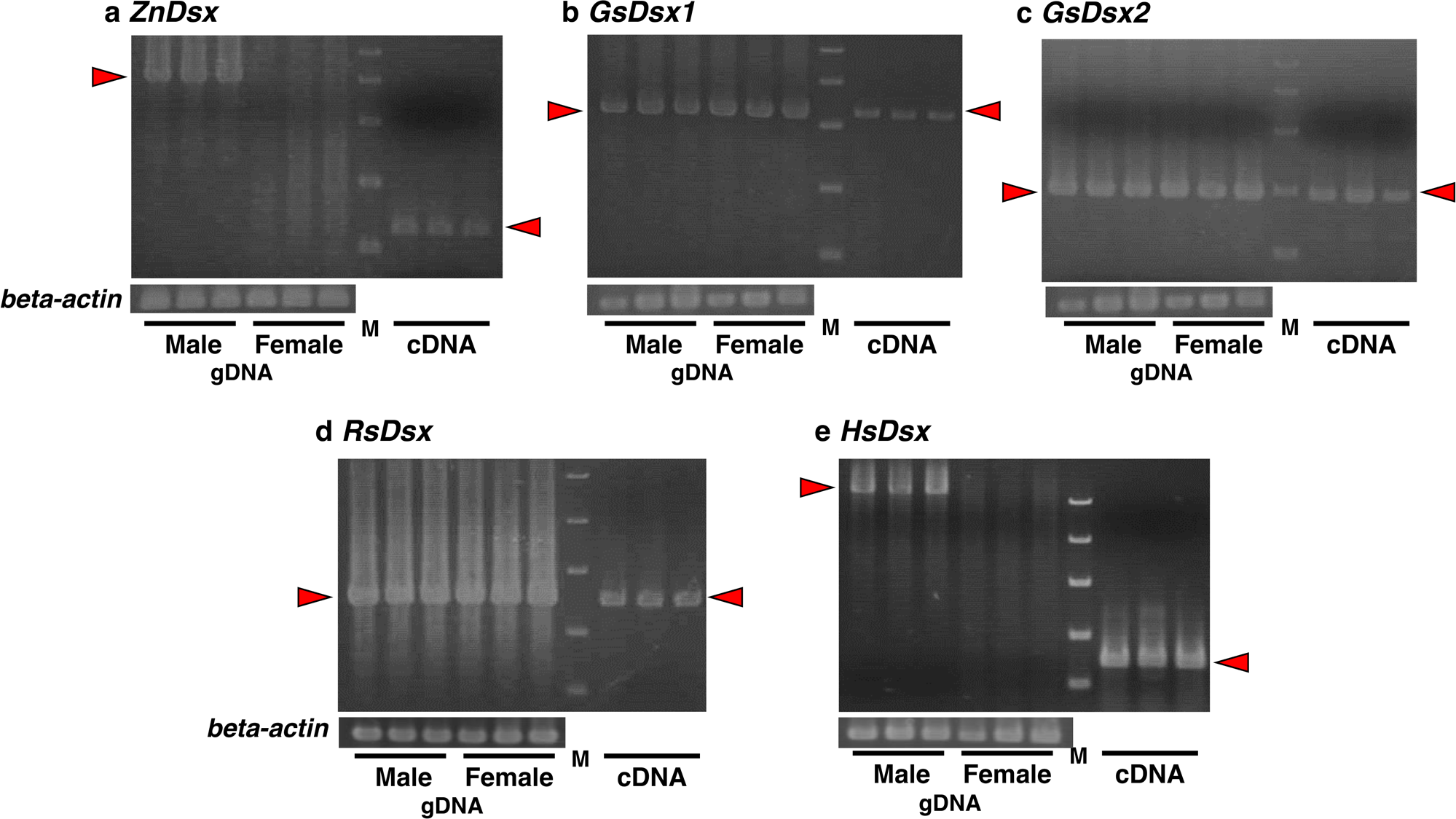
Amplification product sizes using gDNA and cDNA in (a) *Z. nevadensis*, (b-c) *G. satsumensis*, (d) *R. speratus*, and (e) *H. sjostedti*. Left three lanes are amplification products using gDNA extracted from males. Middle three lanes are those using gDNA extracted from females. Right three lanes are those using cDNA derived from nymphs, with equal amounts from each sex. Each lane is the result of a different gDNA or cDNA sample. M indicates the molecular weight of the marker (10000, 4000, 2000, 1000, and 500 bp from the top). Each panel below shows the results of amplification of *beta-actin*, indicating the success of gDNA extraction. The red arrows indicate each band of amplification products.

### 3.3. Synteny analysis

In both *Z. nevadensis* and *H. sjostedti*, gDNA PCR products were obtained only in males, not in females (Figure 4a, e). In *G. satsumensis* and *R. speratus*, however, PCR products were obtained using gDNA extracted from both males and females (Figure 4b-d). Therefore, we compared the genomic location of *Dsx* between *R. speratus* and *Z. nevadensis*, whose genomes have been elucidated (Shigenobu et al., 2022; Terrapon et al., 2014). The synteny of *Dsx* and *prospero* (*pros*) has been confirmed in *B. germanica* and termites including *R. speratus* (Miyazaki et al., 2021; Wexler et al., 2019). We identified *pros* of *Z. nevadensis* (*Znpros*, gene ID: Znev_13276) on the scaffold 502 (gene model Znev_OGSv2.2). *ZnDsx* was not found on the same scaffold, however *UTX homolog* (*ZnUTX*, Znev_13277) was found approximately 25 kb away, adjacent to *Znpros* (Figure S5a). In *R. speratus*, *Rspros*, *RsDsx*, and *UTX homolog* (*RsUTX*, RS012495) were found on the same scaffold in the specified order, with approximately 80-100 kb separating each gene (Figure S5a). Note that the gene model of *R. speratus* (RspeOGS1.0; Shigenobu et al., 2022) was derived from gDNA extracted from female individuals. *Znpros* and *ZnUTX* were amplified from both male and female gDNAs (Figure S5b-c). *Rspros* and *RsUTX* were also amplified in both sexes (Figure S5d-e).

### 3.4. Gene expression analysis

In *R. speratus* and *N. takasagoensis*, *Dsx* is only expressed in males (Miyazaki et al., 2021). The expression levels of *Dsx* relative to those of a most suitable reference gene (Table S3) were examined by real-time quantitative PCR analysis in *Z. nevadensis*, *H. sjostedti*, *O. formosanus*, and *G. satsumensis* (Figure 5a-f). The expression levels of *Dsx* in *C. secundus* were measured using previous RNA-seq data (Harrison et al., 2018) (transcripts per million, TPM; Figure 5g). The results indicated that the expression of *Dsx* was limited to males in *Z. nevadensis*, *H. sjostedti*, and *O. formosanus* (Figure 5a-c), while *Dsx* of *G. satsumensis* and *C. secundus* (Kalotermitidae) was expressed in both males and females (Figure 5d-g). *GsDsx1* and *CsDsx1* were expressed equally in both sexes, while *GsDsx2* and *CsDsx2* showed higher levels in males compared to females (Figure 5d-g).

**Figure 5.**
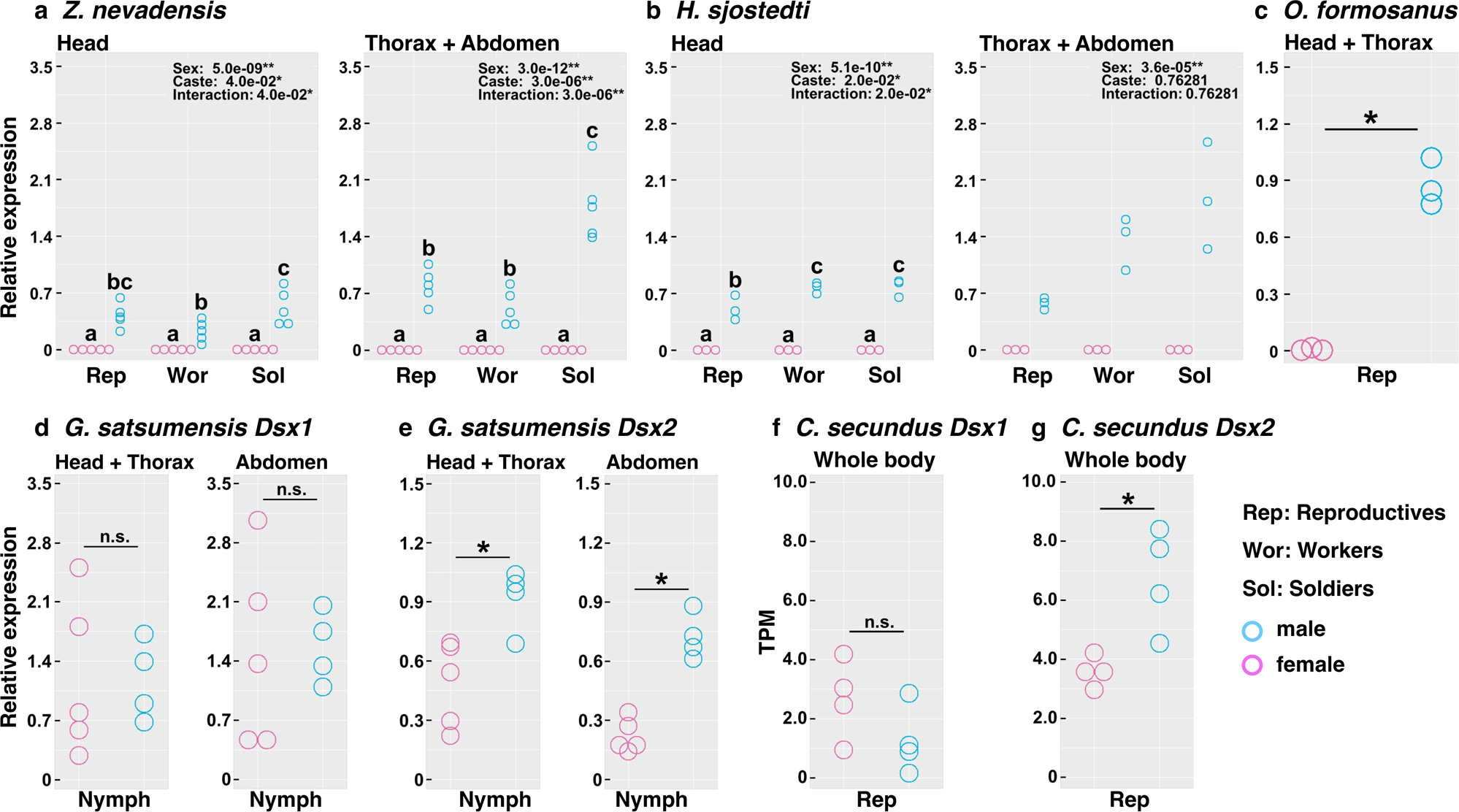
Expression patterns of *Dsx* homologs in (a) *Z. nevadensis*, (b) *H. sjostedti*, (c) *O. formosanus*, (d-e) *G. satsumensis*, and (f-g) *C. secundus*. (a) Real-time quantitative PCR (qPCR) expression levels of *ZnDsx* relative to those of *NADH-dh* among castes in head (left) and thorax and abdomen (right). (b) Real-time qPCR expression levels of *HsDsx* relative to those of *18S* among castes in head (left) and thorax and abdomen (right). (c) Real-time qPCR expression levels of *OfDsx* relative to those of *GAPDH* between female and male reproductives. (d-e) Real-time qPCR expression levels of *GsDsx1* and *GsDsx2* relative to those of *eIF-1A* between female and male nymphs. (f-g) Expression levels of *CsDsx1* and *CsDsx2* (transcripts per million, TPM) between female and male reproductives. (a-b) Statistical results of two-way ANOVA are described in each box (\**P*□<□0.05, \*\**P*□<□0.01). Different letters indicate significant differences (Two-way ANOVA followed by Tukey’s test, *P*□<□0.05). (c-g) Relative *Dsx* expression levels were compared between sexes using the student’s t-test (\**P* < 0.05). Asterisks indicate significant differences, and n.s. means no significant differences.

## 4. Discussion

In this study, we obtained new sequences for *Dsx* homologs in three termite species (Figure S1-4). In a previous study (Miyazaki et al., 2021), *ZnDsx* was not obtained because of the absence of homologous sequences in the available genome sequence data of *Z. nevadensis* (Terrapon et al., 2014). We believe that reanalyzing transcriptome data is highly effective in detecting previously unidentified homologous genes.

### 4.1. Duplication of *Dsx*

In *G. satsumensis*, two *Dsx* homologs (*GsDsx1* and *GsDsx2*) (Figure S3-4) were identified. In a previous study (Miyazaki et al., 2021), two *Dsx* homologs were identified in *C. secundus*, which belongs to the same family as *G. satsumensis* (Kalotermitidae). These two homologs of *C. secundus* are not sex-specific isoforms because they each have a single-exon structure on the different genomic positions (Miyazaki et al., 2021), making it unlikely that alternative splicing is occurring. In contrast, only one copy of *Dsx* was found in each species of the other termite families. Therefore, a duplication event of *Dsx* may have occurred in a common ancestor of the Kalotermitidae.

The duplication of *Dsx* has been reported in some taxa, such as freshwater branchiopod crustaceans (*Daphnia magna*, *D. pulex*, *D. galeata*, and *Ceriodaphnia dubia*). This duplication may have occurred after the divergence of Malacostraca (Toyota et al., 2013). Furthermore, the “*Dsx-like*” genes have been observed only in the Zygentoma, Ephemeroptera, and Phasmatodea, suggesting that duplication occurred early in the evolution of insects (Chikami et al., 2022). Although there are few reports on gene duplication of *Dsx* below the order level, in the southern dogface *Zerene cesonia,* two copies of *Dsx* were found in the genome (Rodriguez-Caro et al., 2021). The novel *Dsx* paralog, probably acquired from a common ancestor of the genus *Zerene*, may be involved in the regulation of sexually dimorphic ultraviolet wing coloration (Rodriguez-Caro et al., 2021). These results support the idea that *Dsx* paralogs observed in the Kalotermitidae (*Dsx1* and *Dsx2*) have different functions associated with sex determination and the formation of sexual dimorphism. In several species belonging to the Kalotermitidae, the existence of gynandromorphs (or intersex individuals) has been reported (Laranjo and Costa-Leonardo, 2017; Miyaguni et al., 2017). These individuals possessed both male and female phenotypes, and have not been observed in other families. Although the molecular mechanism underlying the development of these intermediate traits remain unclear, it would be intriguing to explore whether *Dsx* paralogs with concurrent expression in both sexes (i.e., *GsDsx1* and *CsDsx1*) are involved in these developments. Again it is important to clarify whether there are functional differences between *Dsx* paralogs in the Kalotermitidae.

### 4.2. Diversity of the exon numbers of *Dsx*

From the results of this study and previous research (Miyazaki et al., 2021), a single exon structure of *Dsx* is suggested common within most termite lineages. However, *Dsx* of archotermopsid termites (*ZnDsx* and *HsDsx*) consisted of more than one exon (Figure 4a, e). An increase in the number of exons may have occurred in the common ancestor of the Archotermopsidae. Alternatively, in ancestral termite groups including the Archotermopsidae, multiple exons are present, whereas in derived termite species, *Dsx* may have evolved into single exons. To address this issue, it is necessary to examine the number of exons in the most ancestral termite family, Mastotermitidae (Figure 1).

### 4.3. Diversity of the genomic location of *Dsx*

The termite sex chromosomes are known for male heterogamety (XX-XY) (Luykx, 1990). In species belonging to the Archotermopsidae, the amplification of *Dsx* was confirmed only in male-derived gDNA (Figure 4a, e), suggesting a possible location on the Y-chromosome. Additionally, the synteny of *Dsx* was not conserved in *Z. nevadensis*, compared to other termites and cockroaches with genome information (Figure S5a). Both *Znpros* and *ZnUXT* were amplified from male and female gDNAs (Figure S5b-c). Consequently, it is unlikely that *Znpros* and *ZnUXT* are located on the Y chromosome, but the translocation to the Y-chromosome is occurring in a very limited position around *ZnDsx* and its neighboring genomic regions. We suggest that the translocation of *Dsx* and neighboring regions to the Y-chromosome has occurred in a common ancestor of the Archotermopsidae, and influenced the male-specific expression. However, in the family Rhinotermitidae and Termitidae, where *Dsx* is not believed to be on the Y chromosome, it still exhibits male-specific expression. It is necessary to clarify whether there are differences in the mechanisms regulating the expression of *Dsx* between these two groups.

### 4.4. Diversity of expression patterns of *Dsx*

The expression levels of *RsDsx* were significantly high in male reproductives (thorax and abdomen), but the expression levels were maintained at low levels in male sterile castes (workers and soldiers) (Miyazaki et al., 2021). However, *ZnDsx* and *HsDsx* were highly expressed in all male castes (Figure 5a, b). These differences may be attributed to variations in the caste developmental pathways, which are roughly divided into two types (linear pathway and bifurcated pathway) (Noirot, 1985). In species with a linear pathway (*Z. nevadensis* and *H. sjostedti*), workers have totipotent capabilities for primary reproductives (i.e., all workers have the potential to develop into winged imagoes). In species with a bifurcated pathway (*R. speratus*), the lines for winged imagoes and wingless sterile workers diverged at an early developmental stage. The degree of imaginal-organ developments varies between groups in each pathway; workers and soldiers in species with a linear pathway possess moderately developed reproductive organs. For example, the 4-7th instars (workers) and soldiers have relatively developed ovaries or testes, and both organs gradually develop during larval development in *H. sjostedti* (Oguchi et al., 2016). The development of sexual traits in insects is generally regulated by sex determination pathway including *Dsx* (Verhulst and Van de zande, 2015). Therefore, we suggest that the moderate *Dsx* expression observed in male workers/soldiers, compared to those in kings, of *Z. nevadensis* and *H. sjostedti* is due to the presence of a linear developmental pathway (Figure 5a, b). There is also a possibility that the different genomic locations of *Dsx* (i.e., on the Y chromosome or not) are related to the distinct *Dsx* expression patterns observed in lineages with linear and bifurcated pathways. To verify this possibility, it would be important to compare upstream factors, such as transcription factors that interact with cis-regulatory elements, which regulate the expression of *Dsx* between termite lineages with different developmental pathways.

The precise reason for the high expression of *Dsx* in soldiers of *Z. nevadensis* and *H. sjostedti* is currently unknown. In *N. takasagoensis*, highly expressed levels of *Dsx* have also been demonstrated in soldiers compared to workers (Miyazaki et al., 2021). Investigating the role of *Dsx* in male soldiers of these species will be necessary.

## 5. Conclusion

We examined the gene structure and expression patterns of *Dsx* in several termite species (Figure 6). *Dsx* homologs were newly identified in three species, all of which possessed only the DM domain, as reported for other termites (Miyazaki et al., 2021). However, exon structures and expression patterns of *Dsx* vary among termite families. In the Archotermopsidae, the number of exons has increased (more than one), and translocation to the Y-chromosome might have occurred. In the Kalotermidae, a gene duplication event of *Dsx* likely occurred, resulting in paralogs (*Dsx1* and *Dsx2*) with different expression patterns. All members of the Archotermopsidae and Kalotermitidae exhibit a linear caste developmental pathway. Acquisition of the linear pathway may be associated with *Dsx*-dependent regulatory modifications, originating from the changes in *Dsx* characteristics clarified in this study. Alternatively, the structure and expression of *Dsx* could be simplified to phylogenetically derived groups (Termitidae and some Rhinotermitidae, including *Reticulitermes*). To test these hypotheses, it is necessary to investigate the gene structure and expression patterns of *Dsx* in the most ancestral termite group, Mastotermitidae (comprising only *Mastotermes darwiniensis*; Figure 1), which exhibits a bifurcated caste developmental pathway.

**Figure 6.**
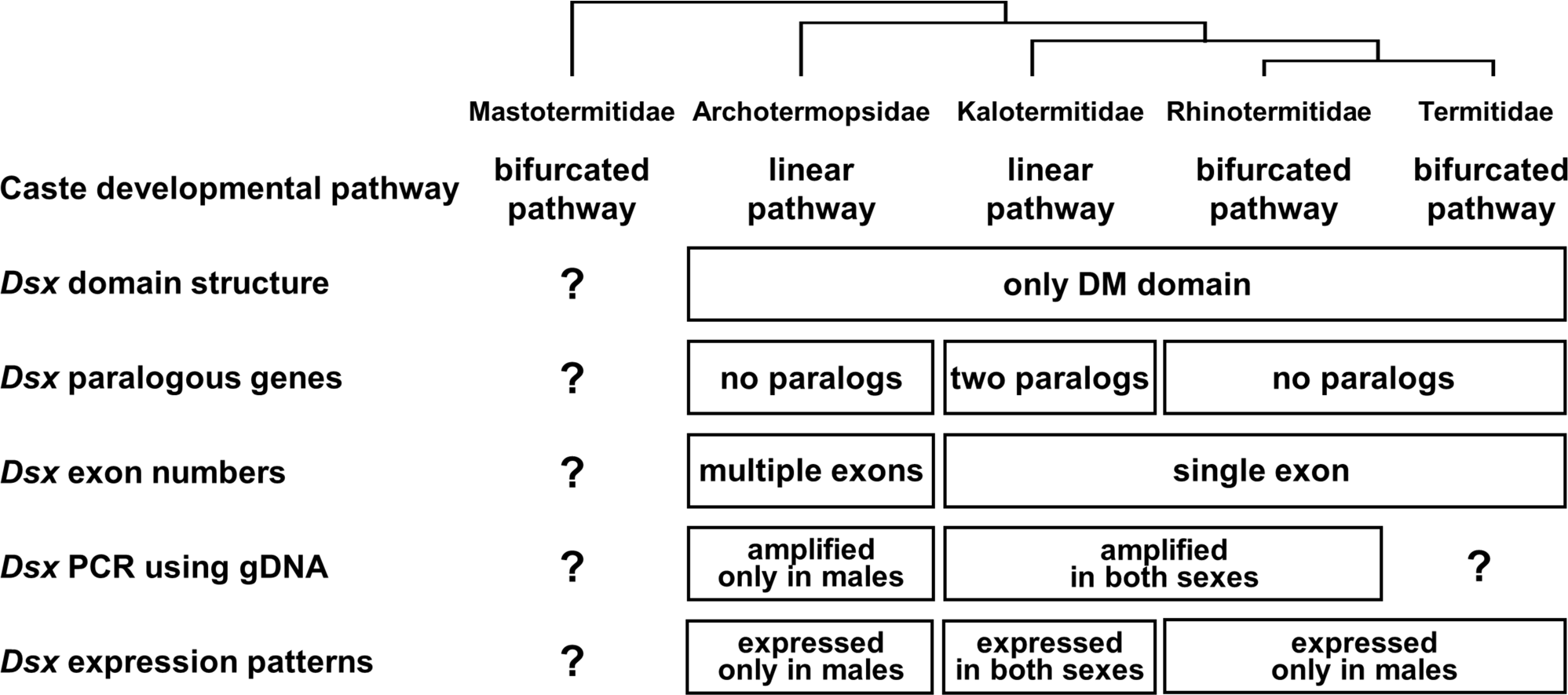
Characteristics of *Dsx* homologs in each termite lineage. A simplified phylogenetic tree and the characteristics of *Dsx* are illustrated. All species within the Archotermopsidae and Kalotermitidae exhibit the linear caste developmental pathway. Remarkable differences in *Dsx* characteristics, such as increased exon numbers (Archotermopsidae) and alterations in expression patterns (Kalotermitidae), are observed in these families.

## Acknowledgements

We are grateful to Masaru Hojo (University of the Ryukyus) for providing us the samples of *Odontotermes formosanus*. We also thank Toru Miura, Kohei Oguchi, Naoto Inui, Soma Chiyoda (University of Tokyo), Takumi Hanada, Mutsuaki Tobo, Ryusei Ashihara, Tousuke Kaneko, and Rina Nakaya (University of Toyama) for their help with field and laboratory work. The experimental analysis was performed using an Applied Biosystems 3500 Serises Genetic Analyzer and QuantStudio 3 Real-Time PCR System at the Division of Instrumental Analysis, University of Toyama. Computational resources for *de novo* assembly were provided by the Data Integration and Analysis Facility, National Institute for Basic Biology.

## Funding

This study was supported in part by the JST SPRING (No. JPMJSP2145), and Scientific Research (Nos. JP21K19293 and JP22H02672 to KM) from the Japan Society for the Promotion of Science.

## Competing interests

We have declared that no competing interests exist.

## Author contributions

K.F. and K.M. designed the study; K.F. and S.M. conducted the experiments; K.F., S.M., and K.M. analysed the data; K.F. and K.M. drafted the manuscript; all authors contributed to the final version of the manuscript.

## Supplementary materials

**Table S1.** Information of OTU used for phylogenetic analysis in this study.

**Table S2.** Primer sequences used in this study.

**Table S3.** Stability values of reference genes in real-time qPCR analysis.

**Figure S1. DNA sequence of *ZnDsx*.** Yellow regions indicate DM domain. A pair of primers (A1) was used for real-time qPCR. A gene-specific primer (B1) was used for RACE PCR analysis and a pair of primers (C1) was used for exon structure analysis.

**Figure S2. DNA sequence of *OfDsx*.** Yellow regions indicate DM domain. A pair of primers (A1) was used for real-time qPCR. A gene-specific primer (B1) was used for RACE PCR analysis.

**Figure S3. DNA sequence of *GsDsx1*.** Yellow regions indicate DM domain. A pair of primers (A1) was used for real-time qPCR analysis. A pair of primers (B1) was used for exon structure analysis.

**Figure S4. DNA sequence of *GsDsx*2.** Yellow regions indicate DM domain. A pair of primers (A1) was used for real-time qPCR. A pair of primers (B1) was used for exon structure analysis.

**Figure S5. The synteny of *Dsx* and the results of PCR amplification of *pros* and *UXT* homologs.** (a) A diagram illustrating the synteny of genes in *Z. nevadensis* and *R. speratus*. *The approximate distance between *Rspros* and *RsDsx* has been reported in previous study (Miyazaki et al. 2021). (b-c) Amplification products of *Znpros* and *ZnUTX* using gDNA extracted from males and females in *Z. nevadensis*. (d-e) Amplification products of *Rspros* and *RsUTX* using gDNA extracted from males and females in *R. speratus*. M indicates the molecular weight of the marker (100-bp intervals from 1,000 bp (top) to 100 bp).

## Notes

### Competing Interest Statement

The authors have declared no competing interest.

