## Supplemental Fig. 1 for "Evolution of the sex-determination gene *Doublesex* within the termite lineage"

ATGGTTCTTCACGTCTGTGAAATGTATCAGAGCAGATCACTGATCTATAGCACTCTCCGATATTGTGCCCGTTG
C1

TGGAAATCATGGATTACGAATGCGTCTGAAAGGGCACAAACATTACTGCTTTTTTCGAGACTGTCCCTGCTAC

AAATGTTCTCAAACAGCGGAGCGCCAAAGAAGTATGGCACGACAAATAGCTGAGCGGCGTGCACAGGCGCT
A1

TGATGAGGACCGCCACGCTAGCAGGTTCAGACCGATTACTAATGGAAACACACTAGTCACCCCCAGAAACTC
A1 B1

CCAACTTGAGTCGTCTGCCGAAGACAAGCCTGTCACAATTTTTCTTTCAGCTGGGAAAAATCAGAATTCGTCC

TCTTCGCACTTTTATAGAGGTGACAGGTCCCTCTCATCTACGTCGACTCCGACGATAAACATCTCACCGCAGC

TGAACTTAACGAGCATAATCTCTGAAGAAAATATGCCGTCTGTGGCACTGTGGGATGCTCCCATATTATTGCAC

AACCAGAGAGGATTGTACTACCAGATTGGTCCTTACTGTCATGACTGCTGCTATATGTCTTCTGCATTATCTAAC

ATCGAGTTCGGTCTGTACCCAACACCACCTGAATTTAACCTACTAAATTTCATAAACTGTTAA
C1

Figure S1
