## Supplemental Fig. 2 for "Evolution of the sex-determination gene *Doublesex* within the termite lineage"

ATGAAGACGATAGGCTCCCGGCCGGATCCAAGGATATCAGTGGCCAGCACTAGCAAGAGGTCACACAAAAAGGACACCGTC

TCCAGGTTGAGGATTAACCGTGCCATACCACAATGTACCCGATGTCTAGCTCATGGCAAATCATTTCCTGTCAAAGGACACAA

ACCTCATTGTAAGCATCGTAACTGTGTTTGTGACACGTGTGTAGTGGCAGAGAAAGCTAGTTGGTATAGAGCGAAGCTGTTG

A1

GCTAAGATGCGCAGTATAAAAAACAAAAAGGCCAGTGAAACAGCTGAAACCGATTCAAACGCAGTAGCGGGTAGTTTACCGT

A1

TTCGTTCATTTCTTTGTAACAGTGCTCAGAGTCTGTCCACCAGTCGCGCGCCAAATGCCAGTTGCAACATAAGCCCGCTGCC

CCAGTCTCAGCATCAGTTAACACGGCGGTGCCAACAATGTTTTGAAAACCGTACGCCATTGAACACTGCGGACACTATAGTTG

B1

AAGAAAGAGTACTGCCATCTATTAGCGGGATAACACAGCCAGACGTGTCCAGCGACTCGCTCTCATCGTCCGGTACCACCAG

CGACAGAAACTACCCAGGCACCTATATTAACATCGACTGGAGCACAACTAAACGACCGCAGCAAAACCAACAATGTCCACTTG

GGTGCAGTCACAGTTTCACAAATTATGAGGCACAGCCAGACAACACCACTCCTATGACAAATTTACTCTCAATGATAGACTGG

B1

AGTAGCTCCCTATCAAGTTCCACCTCCAACACAACAGTGCCAAACGTGCCCAGCAGCACAAATCCATCTCCCAATACAAAGAC

ACAGTACCATCTACGGAGCAAAGCTGAAGCCCCTAGTACGGATTTCCGCTCCATCGACTCTCTATCCTTATCATCTACTTCTGA

AAACAGCAGAACACCGCCAGTTACATCCAGCGACTGGACGAATGTCGCCCTCACAACAATTCCACCGCGGCAGACAGTGAT

GCAACAGCCAGAGCGTTCCTGTGGAAACTGCGAAGCTCTGGCGCAGTGGGAAAGTCTATTTTATGGCTCTTTAAAATTGCAA

TACCCCTTGTATTATCAACATATCAACAGAAGTTAA

Figure S2
