## Supplemental Fig. 3 for "Evolution of the sex-determination gene *Doublesex* within the termite lineage"

ATGGATAAGGAACACAATGAACCGTCTCCAGGGACTTCAGGGTCCACGGCCAGCACTTCAGAAGCCACCGCCAGCAGTTCGGAGGCCACGGGCGGCAGTTACCCAACTTGTGCTAGGTGTCGTAATCATTGGT

B1 A1

TAAAAGTCCGTGTTAAGGGACACAAACGCTTCTGTAAGTACCGAGATTGCACGTGTGACAAATGCTGTCTCGTAGCGAAGAGTCAGAAATACACCGCGCTCCAGACATCGGTGTGGCGCGCACAGGCCCAGGA

A1

CGAGGCCCTCCTTGCCCAGCAAGCCAATCAGGAGGGTGTAGCTGTCACTCGCAAGTTTCCCCCCCAGTTGGATTCGAGATTAAAGGAAAGGATTCGAGACTTCTTGGATTCCACGTCACACGGTACCAAAGGC

GGCAAAAAACGGGTACGTTCTCCATCCGCCCAAAGAACGGATTCCCGTTTAGTGGACAGCAGTAACAACTCCTTGTCTTCTGGGGGCAGAATTCTATCCGCCGCCCCTGCGACGGATTCCCGCTCACTGGATG

AGAGAAGCGACCCGAGATATTCTTCACACAGTTCCACCGCCGGCACAACACAGCAAGCTGCACAGAAATGGAGTGGCTACTCCTTGTGTACCGGAGACCGAGCAATGCCGACAGCCCCAGTCGCCACGTTTCC

TACCGCTCCTGCCGCCGCCGCCGCCGCTCCAAAAATGGATTCCCGAGCAACAGACAGCAGTGTGCTCACCGCTCCAGTCATCTCAGCCGCCCCTGACACGGATTCCCGGGCAGTGGATGGAAGTTGCTATCC

TTTGTCTGCTGGAAGCAGAACTCTGCCCACGGCCCCAGTGACGGACTTTTTGTCACTGTATCCTTCACGTAGTTATGCCACCGGCAAAATATTGCCCATCGCCCCAATCGCTACTCTTCCCACCACCCCTATCGT

CTCTTCACGTCCTGCAGCGAATTTCCGAGCAATAGAGAGTGATTACTCGTTGTCTACCGGTAACAGAGCATTGCTCACCGCCTCCGCCGCCGCTGACACAGATTCCCGAGCAGTGGATGGAGGTTGCTATCCTT

TCTCTACTGGAGGCAGAACTCTGCCCACGGTTCCAGTGACGAACTTTCTCTCAGTCGAGGGGGCCCACAAAGCACTGTATCCTTCACTCGGTTCCACGTCCGGCAAAACACTTACAATCGTCCCAATAGCCACT

CTTCCCACCGCCCCTATCGCCTCTGCCGCCCCTACAGTGGATTCCCGGATAGTAGAGAGCAGTGACTACTCTTTGTCTACCGGTAACGGAGCATTGCTCACGGCCCCAGTCACCTCCGCCGCCCCTGACACAG

ATTCCTGGACAGTGAATGGAAGTTGCTATCCTTTCCCTGCTGGAGGCCGAACTCTCCACACGGTCCCAGTGACGAACTTTCTCTCAGTCGAGGGGTCCCATAAAGCACTGTATCCTTCACTCAGTTCCACGTCC

AGCAAGACATTTCCCACCGCCCCTTCAGAGGATTCCCTCGCAGTAAAGAGGAGCGACTACTCTTTGTCCACCGACAACAGAGCGTTGCTCACCGCCCCAGTCACCTCCGCCCCTGACACTGATTCCCGAACAG

TGGATGGAGGTTGCTATTCTATCTCTGCTGGAGGCAGGACTCTGCCCACGGCCCCAGTGACGGACTTTCTCTCAGTAGAGGGGACCCACAAAGAGCTGTATCCTTCACTCAGCTCCACGTCCGGCAAGACACT

TCCACTTCCAATCGCCCTAACAGACACTCTTTCCACCGCCTCCTTCTCCGCCGCCCCTGCATTGGATTCTCAGGCATTAGAGAGGAGCGACTATTCGTTTTCTACCGGTAACAGAGCATTGCTCACAGCTCCATT

CGCCACCGCAGCCCTTGGCACGGATTCCCGGACAGTGGATAGAGGTTGCTATCCTTTGTCTGCTGGAGGAAGAACTTTGCCTACGGCCCCAGTGACGGATTTTCTCTCTGTGGAAGCGGCCAACAAAGCACTG

TATCCTTCATTCACTTCCACGTCCGGCAAGACACTTCCCATCGTCCCAATAGCCACACTTCCCACCGCCCCAATCTCCTCCACCGCCACTGCTGTGGATTCCCGCTTAATGGAAGGCAGTTCCTATTCTTTGCCT

GCCGGAGGCAGAACACTACCCATTGCCCCATACGGCTTTGCTCCTCCTTGGATGGATTCAAGCCCAATGGAAGGGGGTGGCAGATCACTGCCCACCGCGTTCAGCGTCAAGACAAACCCCTCGCAGTTACCG

CTTTGGTGCGGTCAGGAGTTCCCAATCCGTGGGTCAGTGACCAGTGCCTATAATGATATGTTATTAAATCGGTACTACCAGAACTTGAGCTATTTAACCAAACCGCATTAA

B1

Figure S3
