## Supplemental Fig. 4 for "Evolution of the sex-determination gene *Doublesex* within the termite lineage"

ATGGAGAGGGAAGGCACTCGACTGCTTGCAGGAACTTTCGGGGCCACAGGCAATAGTCAAGGAGCCACTTTCAGCAGTCCGGA

B1

AGCCACAGCCACGAGTTCAAACTTCCCAAACTCTTACCCAACTTGTTCTAGGTGTCGCAATCATTGGTTAAAATCCGCGTGAAAG

GACATAAACGGTACTGTAAGTACCGAGATTGTACGTGTGAGAAATGCTCCCTGGTAGCGGAGGGTCAGAAAATTTCGGCGCTTCG

GATGGCGGTGTGGCGCGCCCAGGCACAGGATGAAGCCCATATTGTCAAAGAAATCAGTCAGCAGCCGGAAAAGGGGGTAACTG

CCACCCACAGTGTACCGTCTGCGTTGGATTCTGGTTTAAAAGAAAGGATTCAGAACTTAATTCGCTCTTTACACACAGTCACCAGC

GGCAGAAAACAGAGCTCCGCCGTATTCTCCCTTGGAGTGGATTGCCGTTCAACTGTGAGTAGTAGCTATTCCTTATCTGCCGGCG

A1

GCACAACTCTGTCCACTACCTTCACCACCCCTGTGACATGTTTCCCTTCATTGGGTGAGAGCATCGACTCTCTTTATTCTCCACAC

A1

AGTTCCACTGCACCACAGGCCACTGCGCTAGTTGTCTCTGGCGCCCCAGTGACAGATTCCCAAGCATTAGTGGGAAGTGGCTAC

CTTCTGTCTACCGGAAACAGAAGACTGACCGCCATCCCTTGGATTGATTCCAACCCAGTGCAGGGCTGCTCCTATTCTATGCCTG

TTGGTAGCAGAATGTTACCCACTGCCTCATTGTCCTCTGACGCCCTTTGGATGGATTCCAGGTCAATTGAGTGGACCGGCTACAC

TGTATCCACCGGCGGTACATCGCTGCCCACCGTGTCCAGCGCCGAAACTAACTCCTCGCAGTTTCCGATTTCGAACAATCAGGTT

ATCCAAAACTGCGGGTCTATGACCAATGTCTGCTATGAGATGTTGTTAAATCAGTACTATCTTAACTTGCGCTATTTAATCAAACCTT

B1

CTTGA

Figure S4
