## Supplementary figures and images for "Evolution of the sex-determination gene *Doublesex* within the termite lineage"

### Supplemental Fig. 5

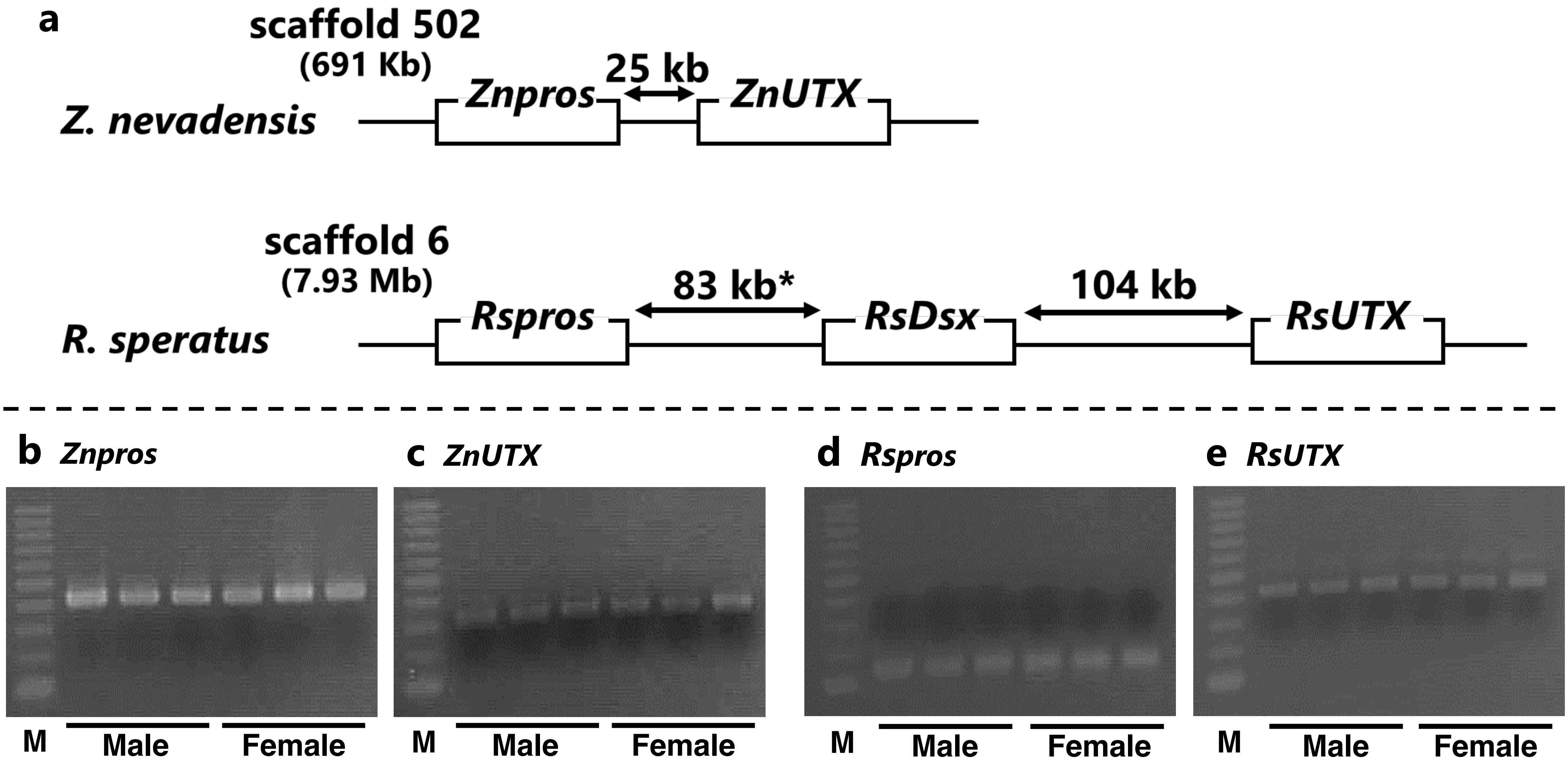
