## Supplemental Table 1 for "Evolution of the sex-determination gene *Doublesex* within the termite lineage"

Table S1. Information of OTU used for phylogenetic analysis in this study.

| Clade | Gene | Family (termites) | Species | Accession number |
|---|---|---|---|---|
| *doublesex* | *doublesex* | Termitidae | *Nasutitermes takasagoensis* | LC635718 |
| | *doublesex* | Termitidae | *Macrotermes natalensis* | Mnat_08109 |
| | *doublesex* | Termitidae | *Odontotermes formosanus* | XXXXX |
| | *doublesex* | Rhinotermitidae | *Reticulitermes speratus* | LC635717 |
| | *doublesex* | Rhinotermitidae | *Coptotermes formosanus* | scaffold506: 427884..429431 |
| | *doublesex1* | Kalotermitidae | *Cryptotermes secundus* | XM_023861307 |
| | *doublesex2* | Kalotermitidae | *Cryptotermes secundus* | XM_023858380 |
| | *doublesex1* | Kalotermitidae | *Gryptotermes satsumensis* | XXXXX |
| | *doublesex2* | Kalotermitidae | *Gryptotermes satsumensis* | XXXXX |
| | *doublesex* | Archotermopsidae | *Zootermopsis nevadensis* | XXXXX |
| | *doublesex* | Archotermopsidae | *Hodotermopsis sjostedti* | LC635719 |
| | *doublesex* | | *Cryptocercus punctulatus* | LC635715 |
| | *doublesex* | | *Blattella germanica* | MK919541 |
| | *doublesex* | | *Drosophila melanogaster* | NM_169202 |
| *Dmrt11* | *Dmrt11* | Termitidae | *Nasutitermes takasagoensis* | TR57579, comp25194 |
| | *Dmrt11* | Termitidae | *Macrotermes natalensis* | MN011923 |
| | *Dmrt11* | Rhinotermitidae | *Reticulitermes speratus* | RS007930 |
| | *Dmrt11* | Rhinotermitidae | *Coptotermes formosanus* | GFG33987 |
| | *Dmrt11* | Kalotermitidae | *Cryptotermes secundus* | XM_023863338 |
| | *Dmrt11* | Kalotermitidae | *Gryptotermes satsumensis* | XXXXX |
| | *Dmrt11* | Archotermopsidae | *Zootermopsis nevadensis* | XM_022076352 |
| | *Dmrt11* | Archotermopsidae | *Hodotermopsis sjostedti* | Hsjo_m.15983, c18070 |
| | *Dmrt11* | | *Cryptocercus punctulatus* | Cpun_m.20374, comp1991 |
| | *Dmrt11* | | *Drosophila melanogaster* | NM_078591 |
| *Dmrt93* | *Dmrt93* | Termitidae | *Nasutitermes takasagoensis* | comp174542 |
| | *Dmrt93* | Termitidae | *Macrotermes natalensis* | Mnat_01812 |
| | *Dmrt93* | Rhinotermitidae | *Reticulitermes speratus* | RS006912 |
| | *Dmrt93* | Rhinotermitidae | *Coptotermes formosanus* | Scaffold9383:409438-409572 |
| | *Dmrt93* | Kalotermitidae | *Cryptotermes secundus* | XM_023863921 |
| | *Dmrt93* | Kalotermitidae | *Gryptotermes satsumensis* | XXXXX |
| | *Dmrt93* | Archotermopsidae | *Zootermopsis nevadensis* | Znev_05388 |
| | *Dmrt93* | Archotermopsidae | *Hodotermopsis sjostedti* | Hsjo_m.51385, c38968 |
| | *Dmrt93* | | *Blattella germanica* | Scaffold353:1209444-1209310 |
| Dmrt99 | *Dmrt99* | Termitidae | *Macrotermes natalensis* | Mnat_08410 |
| | *Dmrt99* | Rhinotermitidae | *Reticulitermes speratus* | RS002870 |
| | *Dmrt99* | Rhinotermitidae | *Coptotermes formosanus* | GFG38119 |
| | *Dmrt99* | Kalotermitidae | *Cryptotermes secundus* | XM_0238500541 |
| | *Dmrt99* | Kalotermitidae | *Gryptotermes satsumensis* | XXXXX |
| | *Dmrt99* | Archotermopsidae | *Zootermopsis nevadensis* | Znev_16235 |
| | *Dmrt99* | | *Drosophila melanogaster* | NM_079704 |
