## Supplemental Table 2 for "Evolution of the sex-determination gene *Doublesex* within the termite lineage"

Table S2. Primer sequences used in this study.

| Gene symbol | Species | Gene ID | Experiment | Forward sequence (5' - 3') | Reverse sequence (5' - 3') | Reference |
|---|---|---|---|---|---|---|
| *OfDsx* | *O. formosanus* | xxxxxx | Dsx search | GTCAAAGGACACAAACGTTCC | GCACTGTTGTGTTGGTGGTG | This study |
| *ZnDsx* | *Z. nevadensis* | xxxxxx | 3' RACE | GTCACCCCCAGAAACTCCCAACTTGA | | This study |
| *OfDsx* | *O. formosanus* | xxxxxx | 3' RACE | GAAAACCGTACGCCATTGAACACTGC | | This study |
| *OfDsx* | *O. formosanus* | xxxxxx | 5' RACE | | GTGGTGTTGTCTGGCTGTGCCTCAT | This study |
| *RsDsx* | *R. speratus* | LC635717 | exon analysis | AATACGACAGACTCCGGGTTCG | GCGCGAATTCAAGTCGTAATGC | This study |
| *ZnDsx* | *Z. nevadensis* | xxxxxx | exon analysis | GGTTCTTCACGTCTGTGAAATG | TTCAGGTGGTGTTGGGTACA | This study |
| *HsDsx* | *H. sjostedti* | xxxxxx | exon analysis | TCGCATCCTAGAATTGCAGTG | TATGGAGCGCCTATGCTTTG | This study |
| *GsDsx1* | *G. satsumensis* | xxxxxx | exon analysis | AGGAACACAATGAACCGTCTCC | TGCGGTTTGGTTAAATAGCTCAAG | This study |
| *GsDsx2* | *G. satsumensis* | xxxxxx | exon analysis | ACTCGACTGCTTGCAGGAACTT | AACATCTCATAGCAGACATTGGTCA | This study |
| *OfDsx* | *O. formosanus* | xxxxxx | exon analysis | GCCGGATCCAAGGATATCAG | CCATAAAATAGACTTTCCCACTGC | This study |
| *beta-actin* | All species | n.a. | exon analysis | GGTCGTACCACCGGTATCGT | CGGATGTCGACGTCGCACTT | This study |
| *ZnDsx* | *Z. nevadensis* | xxxxxx | real-time qPCR | GAAGTATGGCACGACAAATAGCTG | TGGGGGTGACTAGTGTGTTTCC | This study |
| *EF1-alfa* | *Z. nevadensis* | AB915828 | real-time qPCR | GCATGCACTGTTGGCTTTTA | TTCCTCAAATCGGGTTTCAG | Masuoka et al. 2015 |
| *beta-actin* | *Z. nevadensis* | AB915826 | real-time qPCR | AGCGGGAAATCGTCCGTGAC | CAATGGTGATGACCTGCCCAT | Masuoka et al. 2015 |
| *NADH-dh* | *Z. nevadensis* | AB936819 | real-time qPCR | CGGCAAGGAAGCAAATAAAG | CAATGGTGATGACCTGCCCAT | Masuoka et al. 2015 |
| *RS49* | *Z. nevadensis* | KDR21989 | real-time qPCR | CATGCTTCCTACTGGCTTCC | AATTTCGGCACAGAATTTGC | Terrapon et al. 2014 |
| *RPL13a* | *Z. nevadensis* | KDR22610 | real-time qPCR | CACTTCAGAGCACCAAGCAA | ACGTTTCAATGCTGCCTTTC | Terrapon et al. 2014 |
| *RPS18* | *Z. nevadensis* | KDR22651 | real-time qPCR | CTCCGTGAAGACCTGGAGAG | CGTCTTCGTGTGTTGTCCAC | Terrapon et al. 2014 |
| *Znpros* | *Z. nevadensis* | Znev_13276 | real-time qPCR | CACATGAACCCGTTCTGCATGT | GTCACCTTATGGCGCTTCTTCT | This study |
| *ZnUTX* | *Z. nevadensis* | Znev_13277 | real-time qPCR | TGTTGAGAAGCGAGTGAATCTCTT | GGTGCAGCCCTCCTAGTACAA | This study |
| *HsDsx* | *H. sjostedti* | LC635719 | real-time qPCR | TGTCGCCGGTGTCTAAATCA | CCGCTATGCGACAGCATTTT | This study |
| *18S rRNA* | *H. sjostedti* | n.a. | real-time qPCR | ACGTAATCAACGCGAGCTTAT | CTTGCAATTGTTCCCCATGA | Oguchi et al. 2022 |
| *EF1-alfa* | *H. sjostedti* | n.a. | real-time qPCR | TGCTGGTACTGGCGAGTTTG | GCGCATGCTCACGTGTCT | Oguchi et al. 2022 |
| *GAPDH* | *H. sjostedti* | n.a. | real-time qPCR | TGAGTCATACAACCCATCATTGAAG | TTAGCAAGAGGTGCCAAGCA | Oguchi et al. 2022 |
| *RP49* | *H. sjostedti* | n.a. | real-time qPCR | TCAGTACCTCATGCCTACAATTGG | GGAAGCCAGTAGGAAGCATGTG | Oguchi et al. 2022 |
| *OfDsx* | *O. formosanus* | xxxxxx | real-time qPCR | TGACACGTGTGTAGTGGCAGA | GCTGTTTCACTGGCCTTTTTG | This study |
| *beta-actin* | *O. formosanus* | n.a. | real-time qPCR | CTGGAGAAGTCATACGAGTTG | AGAAGGAAGGCTGGAACA | Gao et al. 2020 |
| *NADH-dh* | *O. formosanus* | n.a. | real-time qPCR | TTGGTGAGATTGGTCTGCTG | ACAATGTGTAAGCCGCACTA | Gao et al. 2020 |
| *GAPDH* | *O. formosanus* | n.a. | real-time qPCR | TCACTGCAACGCAGAAGACT | ATTGGAAACTGGCACACGGA | Xu et al. 2021 |
| *GsDsx1* | *G. satsumensis* | | real-time qPCR | GCGGCAGTTACCCAACTTGT | TTGTCACACGTGCAATCTCG | This study |
| *GsDsx2* | *G. satsumensis* | | real-time qPCR | CAGAGCTCCGCCGTATTCTC | GGGGTGGTGAAGGTAGTGGA | This study |
| *EF1-alfa* | *G. satsumensis* | n.a. | real-time qPCR | TTTGCTGTGCGTGACATGAG | CCTTCTCAGCAGCCTTCGTT | This study |
| *beta-actin* | *G. satsumensis* | n.a. | real-time qPCR | GCTATGTCGCCCTGGACTTC | TGACCTGACCATCAGGCAAC | This study |
| *NADH-dh* | *G. satsumensis* | n.a. | real-time qPCR | GCTGGGCTAGGTGCTAATTTTG | GCCAAACCTACAGACACAGCA | This study |
| *EIF-1* | *G. satsumensis* | n.a. | real-time qPCR | TTGTGTCATATCCGTGGCAAG | CATCTGCCTTTGCATCTTGG | This study |
| *RPL13a* | *G. satsumensis* | n.a. | real-time qPCR | CCACCAACCTCTTCCTCCTG | GGGTATGCTTCCCCACAAGA | This study |
| *RPS18* | *G. satsumensis* | n.a. | real-time qPCR | TGGGAAGCGGAAGGTAATGT | TCCAGCCCTTTTGTCCAGAT | This study |
