## Supplemental Table 3 for "Evolution of the sex-determination gene *Doublesex* within the termite lineage"

Table S3. Stability values of reference genes in qPCR analysis.

*Z. nevadensis*

| Gene symbol | Accession no | Stability value | |
|---|---|---|---|
| | | GeNorm | NormFinder |
| *EF1-alfa* | AB915828 | 0.550 | 0.139 |
| *beta-actin* | AB915826 | 0.876 | 0.560 |
| *NADH-dh* [1] | AB936819 | 0.510 | 0.120 |
| *RS49* | KDR21989 | 0.552 | 0.176 |
| *RPL13a* | KDR22610 | 0.773 | 0.465 |
| *RPS18* | KDR22651 | 0.658 | 0.331 |

[1] *NADH-dh* was selected by GeNorm and NormFinder due to the lowest stability values among six genes analyzed.

*H. sjostedti*

| Gene symbol | Accession no | Stability value | |
|---|---|---|---|
| | | GeNorm | NormFinder |
| *18S* [2] | n.a. | 0.310 | 0.048 |
| *Ef-1* | n.a. | 0.363 | 0.196 |
| *gapdh* | n.a. | 0.329 | 0.121 |
| *RP39* | n.a. | 0.449 | 0.284 |

[2] *18S* was selected by GeNorm and NormFinder due to the lowest stability values among four genes analyzed.

*O. formosanus*

| Gene symbol | Accession no | Stability value | |
|---|---|---|---|
| | | GeNorm | NormFinder |
| *b-actin* | n.a. | 0.860 | 0.585 |
| *NADH* | n.a. | 0.611 | 0.307 |
| *GAPDH* [3] | n.a. | 0.518 | 0.094 |

[3] *GAPDH* was selected by GeNorm and NormFinder due to the lowest stability values among three genes analyzed.

*G. satsumensis*

| Gene symbol | Accession no | Stability value | |
|---|---|---|---|
| | | GeNorm | NormFinder |
| *b-actin* | n.a. | 1.392 | 0.943 |
| *NADH* | n.a. | 0.737 | 0.376 |
| *EF1* | n.a. | 0.639 | 0.278 |
| *RPL13a* | n.a. | 0.876 | 0.359 |
| *eIF1A* [4] | n.a. | 0.603 | 0.084 |
| *RPS18* | n.a. | 0.740 | 0.399 |

[4] *eIF-1A* was selected by GeNorm and NormFinder due to the lowest stability values among six genes analyzed.
